# Targeting glioblastoma tumor hyaluronan to enhance therapeutic interventions that regulate metabolic cell properties

**DOI:** 10.1101/2024.01.05.574065

**Authors:** Edward R Neves, Achal Anand, Joseph Mueller, Roddel A Remy, Hui Xu, Kim A Selting, Jann N. Sarkaria, Brendan AC Harley, Sara Pedron-Haba

**Affiliations:** Department of Chemical and Biomolecular Engineering; Carl R Woese Institute for Genomic Biology; Carle Illinois College of Medicine; Materials Research Laboratory; Cancer Center at Illinois; College of Veterinary Medicine, University of Illinois Urbana-Champaign, Urbana, IL 61801, USA; Department of Radiation Oncology, Mayo Clinic, Rochester, MN 55905, USA

**Keywords:** glioblastoma, hydrogel, tumor microenvironment, hyaluronan metabolism, engineered disease models

## Abstract

Despite extensive advances in cancer research, glioblastoma (GBM) still remains a very locally invasive and thus challenging tumor to treat, with a poor median survival. Tumor cells remodel their microenvironment and utilize extracellular matrix to promote invasion and therapeutic resistance. We aim here to determine how GBM cells exploit hyaluronan (HA) to maintain proliferation using ligand-receptor dependent and ligand-receptor independent signaling. We use tissue engineering approaches to recreate the three-dimensional tumor microenvironment in vitro, then analyze shifts in metabolism, hyaluronan secretion, HA molecular weight distribution, as well as hyaluronan synthetic enzymes (HAS) and hyaluronidases (HYAL) activity in an array of patient derived xenograft GBM cells. We reveal that endogenous HA plays a role in mitochondrial respiration and cell proliferation in a tumor subtype dependent manner. We propose a tumor specific combination treatment of HYAL and HAS inhibitors to disrupt the HA stabilizing role in GBM cells. Taken together, these data shed light on the dual metabolic and ligand - dependent signaling roles of hyaluronan in glioblastoma.

**Significance:** The control of aberrant hyaluronan metabolism in the tumor microenvironment can improve the efficacy of current treatments. Bioengineered preclinical models demonstrate potential to predict, stratify and accelerate the development of cancer treatments.

## Introduction

Hyaluronan (HA) is the main constituent of the brain extracellular matrix and is an essential component of the tumor microenvironment. The biosynthesis and catabolism of hyaluronan has multiple roles in tissue architecture and cell signaling. Dysregulation of these processes is crucial in pathological processes such as cancer, inflammation or tissue remodeling. The role of HA is believed to be strongly dependent on its size (molecular weight), location, and cell receptor density and activity (1). The accumulation of HA, its receptors and enzymes are linked to both local invasion and systemic metastasis and are often associated with poor prognosis for multiple tumor types; various therapeutic strategies have thus addressed hyaluronan cell interactions and metabolism (2–4).

Due to the widely distributed presence in biological tissues and the large array of biological signals, HA has been increasingly used in the design of biomaterial platforms as disease models (5,6). In many tumors, the dysfunctional biosynthesis and signaling of HA involves more threatening cancer progression and therapeutic resistance (7–9). The control of hyaluronan biosynthesis and degradation may become essential in analyzing the role of HA in tumor biology and predicting the functionality of the multiple molecular weight fragments (10). HA biological functions depend, in part, on its molecular weight (MW): oligo-HA (< 3 kDa), low molecular weight HA (LMW, 3–10 kDa), medium molecular weight HA (MMW HA, 10-250 kDa), and high molecular weight HA (HMW, > 250 kDa). The distribution of MWs of HA are determined by the levels of hyaluronan synthetic enzymes (HAS) and hyaluronidases (HYAL). HA is produced intracellularly by HA synthases (HAS1-3): HAS2 synthesizes HMW HA chains, while HAS3 produces shorter polymers of LMW HA (11). HA polymers can also be catabolized endogenously by hyaluronidases (HYAL1-3) which hydrolyze the β-(1–4)-hexosaminidic bond, leading to low molecular weight HA and HA oligosaccharides (o-HA) (12).

The fabrication of controlled microenvironments provided by three dimensional (3D) models helps to elucidate the role of HA signaling in GBM tumors. We exploit here the advantages of 3D in vitro models to understand the important role of hyaluronan in the local behavior of GBM. The dynamic GBM tumor microenvironment leads to the implementation of different strategies to control its progression. We have previously studied tumor progression and widely characterized therapeutic resistance in hyaluronan rich environments (13,14). We have found that signaling from endogenously or exogenously produced hyaluronan is essential in tumor biology, increasing cell survival through the PI3K pathway and STAT3 activation (14). Based on these findings we seek to unwind the continuous HA synthesis and fragmentation influence on tumor growth. A more precise understanding of HA metabolism and the specific effects of the molecular weights of HA on cellular signaling and functionality can provide new perspectives for future druggable targets (15). The development of effective tools that allow the understanding of HA metabolism and signaling, as a microenvironmental factor that contributes to cancer progression and therapeutic resistance, will lead to new therapeutic interventions.

Hyaluronan accumulation in the tumor microenvironment preserves proliferation of GBM cells by different mechanisms after treatment (16,17). Ongoing research suggests that HA might be consumed by mitochondria and used as an alternative bypass energy cycle in solid tumors (18,19). We hypothesize that protective effects of hyaluronan on GBM tumor cells are due to the turnover of specific HA fragments to preserve proliferation and mitochondrial function. Here we adapt a series of gelatin-based hydrogels to investigate the role of hyaluronan in GBM tumor biology and its potential therapeutic targets. We provide a strategy to predict and stratify tumors in order to achieve more efficient combinatorial treatments targeting endogenous hyaluronan production. We show that HA fragments can enhance tumor metabolism and growth through receptor-signaling dependent pathways and receptor-signaling independent pathways. This work emphasizes the potential of these preclinical models to predict and accelerate cancer treatments.

## Materials and Methods

### Cell laden hydrogel fabrication

Gelatin methacrylamide (GelMA) was prepared as described previously (14). We produced 5wt% GelMA scaffolds by UV photopolymerization in the presence of 0.1 wt% LAP (Lithium phenyl-2,4,6-trimethylbenzoylphosphinate) as a photoinitiator. Prepolymer solution was pipetted into Teflon molds (5 mm diameter x 1.5 mm thick) and exposed to 10 mW/cm^2^ UV light (AccuCure Spot, LED 365nm, Digital Light Lab) for 30s. Patient-derived xenograft cells GBM6, GBM8, GBM10, GBM12, GBM34 and GBM44 (Mayo Clinic, Rochester MN) were cultured within these hydrogels at a concentration of 5 million cells/ml. Some relevant GBM characteristics are shown in **Table 1**. Two biological and three technical replicates were performed. Young modulus of these hydrogels is 2.5 ± 0.3 kPa, as measured by compression testing.

**Table 1.** Selected characteristics of patient-derived xenograft cells.

| GBM | Median survival<br>(days) | +EGFR | Erlotinib | Sex | Molecular<br>subtype | MGMT |
| --- | --- | --- | --- | --- | --- | --- |
| 10 R | 26 | NO | resistant | M | M | U |
| 12 | 24 | YES | sensitive | M | M | M |
| 6 | 28 | YES/vIII | resistant | M | C | U |
| 8 | 89 | YES | resistant | F | C | M |
| 44 | 54 | NO | resistant | F | M | U |
| 34 | 75 | YES | resistant | F | P | U |

All IDH1 and IDH2 wild type. R (Recurrent), M (Mesenchymal), C (Classical), P (Proneural), U (Unmethylated), M (Methylated)

### Cell culture and drug treatments

Patient-derived xenograft GBM cells from Mayo Clinic (Rochester, MN) (**Table 1**) were cultured in DMEM medium supplemented with 10% FBS and penicillin/streptomycin (100 U/ml and 100 μg/ml). Cells were cultured at 37°C in a 5% CO2 environment. For hydrogel cultures, patient-derived cells were incubated on an orbital shaker in low adhesion well plates containing standard culture media for 48 hours. After providing 48 hours to stabilize within the hydrogels, cells were exposed to 10 μM erlotinib (Erl, Biovision Inc., in 0.1% v/v DMSO), 0.5 mM 4-Methylumbelliferone (4-MU, Selleckchem, in 0.1% v/v DMSO) and 0.1 mM Glycyrrhizin (Sigma-Aldrich) added to the culture media for 3 days (0.1% v/v DMSO and DMEM were used as a controls). Cell metabolic activity was evaluated by MTT colorimetric assay (CyQUANT, ThermoFisher Scientific). Hyaluronic acid <10 kDa (Lifecore Biomedical HA5K) was used as low molecular weight HA (LMWHA) in media (0.25 wt%).

### Protein immunoblotting

Analysis of protein expression was determined from cell-hydrogel specimens first lysed using RIPA buffer on ice for 30 min. After lysis and determination of protein concentration, samples were separated on a 10% polyacrylamide precast electrophoresis gel (Biorad #456-1096), and transferred onto a nitrocellulose membrane (Amersham Protan, GE Healthcare, #10600012). Proteins were visualized by ECL western blotting detection reagents (Pierce, ThermoFisher Scientific), according to manufacturer’s instruction. The following primary antibodies were used (all rabbit): Vinculin (E1E9V) XP^®^ Rabbit mAb (#13901) and β-actin (#4967S) from Cell Signaling, Danvers, MA; HAS1 (ab128321), HAS2 (ab131364), HAS3 (ab138541), HYAL1 (ab85375), and HYAL2 (ab68608) from Abcam Waltham, MA. Primary antibodies were then labeled with goat-anti-rabbit IgG conjugated to horseradish peroxidase (Cell Signaling). Blocking solution was 5wt% non-fat dry milk in TBST. Imaging signal was visualized using imaging kits (SuperSignal™ West Pico PLUS Chemiluminescent Substrate or SuperSignal™ West Femto Maximum Sensitivity Substrate, Sigma-Aldrich) via an Image Quant LAS 4010 chemiluminescence imager (GE Healthcare). Band intensities were quantified using ImageJ (RRID:SCR_003070) and normalized to vinculin or β-actin expression.

### Mitochondrial Stress Test

The mitochondrial stress test assay was performed using a Seahorse XFe96 analyzer (Agilent Technology, Santa Clara, CA) in the Tumor Engineering and Phenotyping Shared Resource of the Cancer Center at Illinois. Hydrogels laden with GBM34, GBM12 and GBM10 were prepared in small molds to hold 20 μl and polymerized as previously described. GBM cells within hydrogels were seeded at a density of 1 × 10^5^ cells/well in 96-well spheroid microplates (Agilent, catalog number 102959-100). In the Mito Stress Test, on the day of the assay the growth medium was replaced with assay medium (Agilent) as instructed. Cells were incubated in a non-CO2 incubator for 1 h before initiation of the test. The assay medium for the standard Mito Stress Test was Seahorse XF base medium supplemented with 10 mM glucose, 1 mM pyruvate and 2 mM glutamine. The final concentrations of specific chemicals were optimized to be oligomycin (Sigma-Aldrich) at 20 μM, carbonyl cyanide-4-(trifluoromethoxy) phenylhydrazone (FCCP) (Sigma-Aldrich) at 7.5 μM, and antimycin A/rotenone (Sigma-Aldrich) at 5 uM/10 μM. The assay protocol was modified (for GBM34) from the standard protocol by inserting an extra step after each injection of the reagents. The extra step includes 2 cycles of 3-min mix and 7-min wait. This additional assay, adding and extra step, corroborated that diffusion was not hindered during the stress test.

### Immunofluorescent staining

Hydrogels were fixed in Image-iT Fixative Solution (Invitrogen) for 15 min followed by three PBS washes. Cells were permeabilized in a 0.1% Triton-X-100 solution in PBS for 15 min at room temperature followed by three five-minute PBS washes. Samples were blocked for 2 h in a 2% bovine serum albumin (BSA) solution in PBS and subsequently incubated in Hyaluronic Acid Binding Protein, Bovine Nasal Cartilage, Biotinylated (1:100 HABP, Sigma-Millipore) solution diluted in a 2% BSA solution overnight at 4°C. Following five five-minute PBS washes, samples were incubated in Streptavidin, Alexa Fluor™ 488 conjugate (1:500 Thermo Fisher), in a 2% BSA solution overnight at 4 °C. After five five-minute PBS washes, samples were incubated for 30 min at room temperature in a 1:1000 dilution of DAPI (Thermo Fisher) in PBS. Samples were washed with PBS and stored at 4 °C until imaged using confocal microscope Zeiss LSM710. Representative images are shown of n=3 hydrogels.

### Hyaluronic acid isolation and Size Exclusion Chromatography (SEC)

The HA isolation from media samples and the molecular weight (MW) distribution analysis was assessed following a protocol from Cleveland Clinic (Hyaluronan size analysis by agarose gel electrophoresis, http://pegnac.sdsc.edu/cleveland-clinic/protocols/). Briefly, after consecutive digestion-precipitation steps, HA was extracted from media samples and lyophilized. Half of the sample was completely digested with Hyaluronidase to serve as a reference. Media samples without cells were used as background controls. Samples were eluted through a diethylaminoethyl (DEAE)-Sephadex columns (Sigma) to remove sulfated GAGs (20). The remaining sample was used for analysis by SEC. MW distribution was measured using a Tosoh EcoSEC Model HLC-8320 GPC equipped with two Tosoh TSKgel Alpha-M columns connected in series. A built-in, dual flow refractive index (RI) detector was used for MW distribution analysis. The aqueous mobile phase consisted of 0.1 M NaNO3 and 0.02% NaN3. The flow rate was set to 0.6 mL/min at 35 °C. Samples were dissolved with the mobile phase and filtered through a 0.45 µm nylon syringe filter before analysis. Pullulan standards (Polymer Standards Service) were used for conventional molecular weight calibration.

### Radiation

Cells were treated with 8 Gy in a single dose for each experiment at 600 cGy/minute using a Varian® TrueBeam® linear accelerator. The Dmax for our 6 MV beam is 1.5 cm and therefore cells in plates were positioned in a field that provided at least 1 cm margin around all irradiated plates on top of 1.5 cm solid water to allow full build up. Radiation was delivered from beneath the treatment couch to allow the beam to pass through the solid water before reaching the cells in culture.

### Ethics statement

The xenografts used in this study were established with tumor tissue from patients undergoing surgical treatment at the Mayo Clinic, Rochester, Minnesota. The Mayo Clinic Institutional Review Board approved these studies and only samples from patients who had provided prior consent for use of their tissues in research were included. All xenograft therapy evaluations were done using an orthotopic tumor model for glioblastoma on a protocol approved by the Mayo Institutional Animal Care and Use Committee.

### Statistical analysis

All analyses were performed using a one-way analysis of variance (ANOVA) followed by Tukey’s HSD post-hoc test. Significance level was set at p < 0.05 or p < 0.01. At least n = 3 samples were examined for cell proliferation assays and at least n = 3 samples were examined for Western Blot analysis. Error was reported in figures as the standard deviation unless otherwise noted.

The data generated in this study are available upon request from the corresponding author.

## Results and Discussion

The glioblastoma tumor microenvironment is a highly complex and interconnected ecosystem that is composed by glioma stem cells, extracellular matrix (ECM), immune cells, neural networks, and vascular system. The interactions between all these components have been shown to be essential in cancer progression and therapeutic response (21). An improved understanding of the GBM tumor microenvironment may provide additional targets to help stratify patients, predict therapeutic outcomes, or enhance the efficacy of current therapeutic approaches. In this study, we focus on developing hydrogel models to investigate the role of hyaluronan-cell interactions on subsequent signaling, matrix biosynthesis, and cell metabolism.

### Hyaluronan regulates tumor progression and therapeutic response

We have previously shown that ECM-hyaluronic acid interactions influence the efficacy of the tyrosine kinase inhibitor erlotinib, that HA secreted levels are increased in more aggressive GBM types, and that blockade of receptor CD44 plays an essential function in GBM tumor maintenance (6,13). We also showed that α-CD44 affects cell invasion and proliferation (14). It has also been widely described in literature that small fragments of HA (3-5 kDa) have specific effects in cell signaling, through CD44 and EGFR interactions (22), highlighting the different roles of HA secreted by glioma cells and the involvement of the different synthases and hyaluronidases. The more exhaustive study of these processes that is reported here will provide valuable insights into the regulation of the tumor microenvironment (**Figure 1A**).

**Figure 1.**
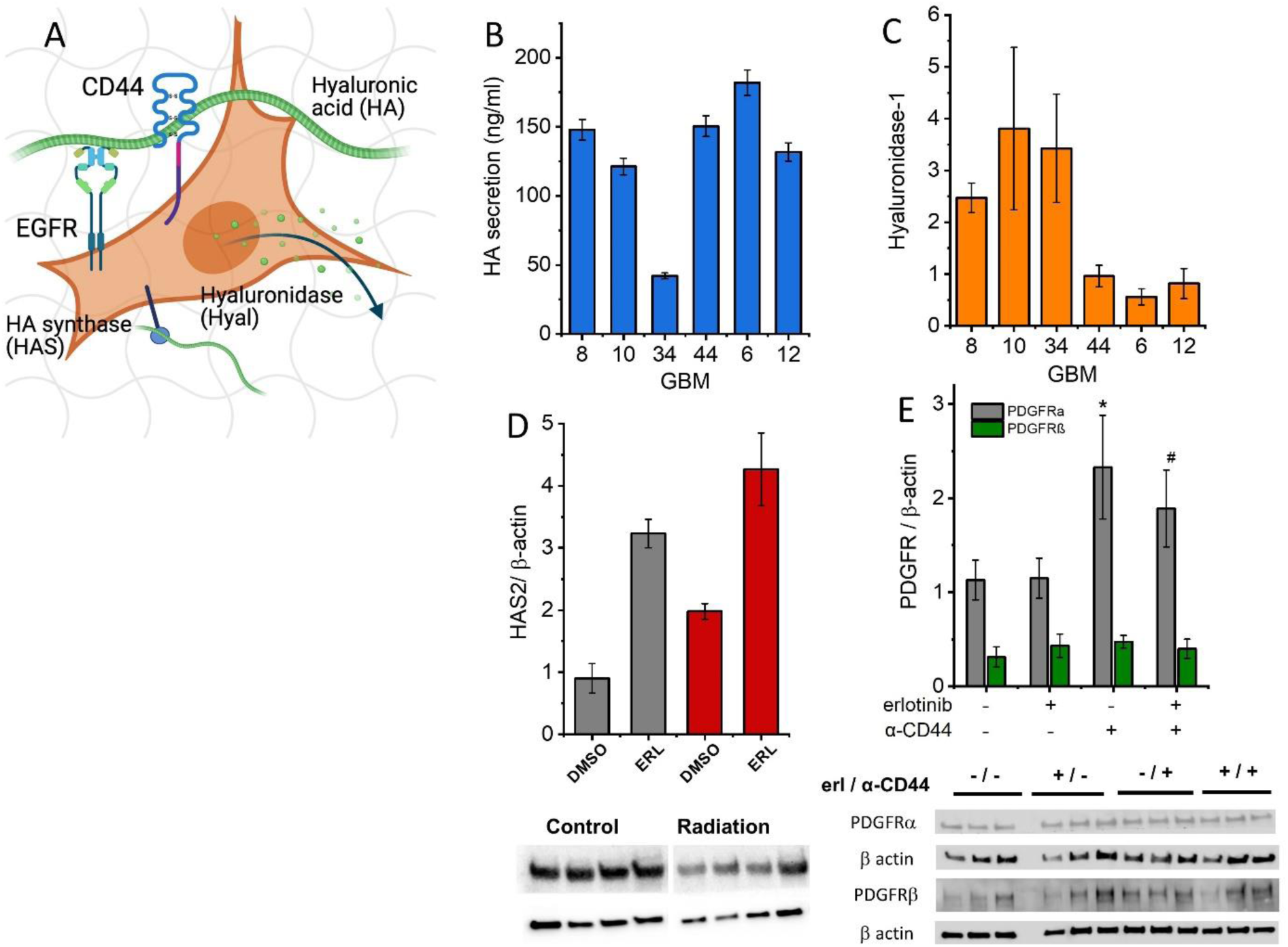
Therapeutic approaches modify HA biosynthesis and signaling pathways. (A) Extracellular matrix and endogenous ECM components regulate tumor cell progression and therapeutic response using different mechanisms of cellular machinery. Biosynthesis and degradation are dependent on tumor subtype. (B) HA concentration using ELISA (Hyaluronan Quantikine ELISA Kit #DHYAL0, R&D Systems), (C) Expression of hyaluronidase-1 detected by Western Blot, vinculin is used as loading control, (D) Western Blot GBM12 HAS2 expression after radiation (8 Gy) and erlotinib treatments, β-actin is used as loading control, (E) Western Blot expression of PDGFR in GBM10 cells within gelatin-HA matrices, exposed to erlotinib and anti-CD44. *# p<0.01, compared to control.

We have carried out a comprehensive study of six GBM xenograft lines derived from patients (GBM6, GBM8, GBM10, GBM12, GBM34, GBM44) that represent a range of different phenotypes and tumor subtypes (**Table 1**). We observe that secretion of HA has great variability depending on the tumor subtype (**Figure 1B**). Hyaluronan is highly expressed in glioblastoma tumors and degradation of HA has demonstrated positive outcomes in combination with antitumor therapies (23). Additionally, tumor cell-derived hyaluronidase-1 (HYAL1) can degrade hyaluronic acid into pro-inflammatory fragments that may support tumor progression. Although HYAL1 has been proven as a decisive factor in tumor progression in different cancers, it has not yet been studied as a therapeutic target (24). We show here that the expression of HYAL1 varies greatly among the different tumors (**Figure 1C**), so inhibiting this enzyme with Glycyrrhizin (GL) could selectively influence cell survival in some patients.

Human hyaluronan is synthesized by three membrane HA synthases (HAS1, HAS2 and HAS3). HAS1 is the only one that occupies primarily cytoplasmic space leading to intracellular HA. HAS1 and HAS2 secrete high molecular weight hyaluronan while HAS3 has been associated to shorter chains (8). The overexpression of HAS and the accumulation of HA in cancer cells, the cancer-surrounding stroma, and ECM has been associated with worse prognoses and treatment resistance. Our results also suggest that GBM cells respond to radiation by increasing production of HA through upregulation of HAS2. We observed here that radiation treated (single 8 Gy fraction) cells have a higher expression of HAS2 in GBM12 cells (EGFR+, erlotinib sensitive) and we found that radiation and erlotinib treatment modify HA biosynthesis and signaling pathways (**Figure 1D**). Radiation is known to affect the HA tumor microenvironment via NF-kB activation (17), increasing invasion, and HAS2 knockdown has been shown to enhance radiation-induced DNA damage in cancer cells (25). The ability to control GBM-HA interaction may provide new avenues to sustain radiation-induced DNA damage.

Moreover, studies show that EGFR-vIII glioblastoma tumors become resistant to EGFR inhibitors by transcriptionally de-repressing PDGFRβ, and therefore opens the path for combinatorial inhibitors in these tumor subtypes (26). PDGFRα amplification has been associated with glioma growth and low survival rates, and the functional interaction with other cell surface tyrosine kinase receptors that present abnormalities in GBM tumors (27–29). We observed here that PDGFRα, but not PDGFRβ, is overexpressed in erlotinib resistant GBM10 cells (EGFR wild type, non-amplified) treated with α-CD44 when HA is present in the matrix (**Figure 1E, Figure S1**). α-CD44 is an inhibitor of extracellular signaling of HA; these results support an essential role of HA in cell maintenance.

### Hyaluronan selectively rescues metabolic activity of glioblastoma cells after treatment with inhibitors that target HA expression

GBM laden hydrogels were exposed to 4-MU and erlotinib (Erl) and the combination of both for 3 days. We have previously described the effect of Erl and matrix-bound HA in these GBM models. The use of HAS inhibitors such as 4-Methylumbelliferone (4-MU) have been reported as tumor growth suppressors (30,31) and inhibit the synthesis of HA depleting GlcA cytoplasmic levels. Using a MTT assay, our results showed that 4-MU is more efficient at reducing cell proliferation than erlotinib. The combination of both inhibitors only showed synergetic effect in GBM34 (**Figure 2**).

**Figure 2.**
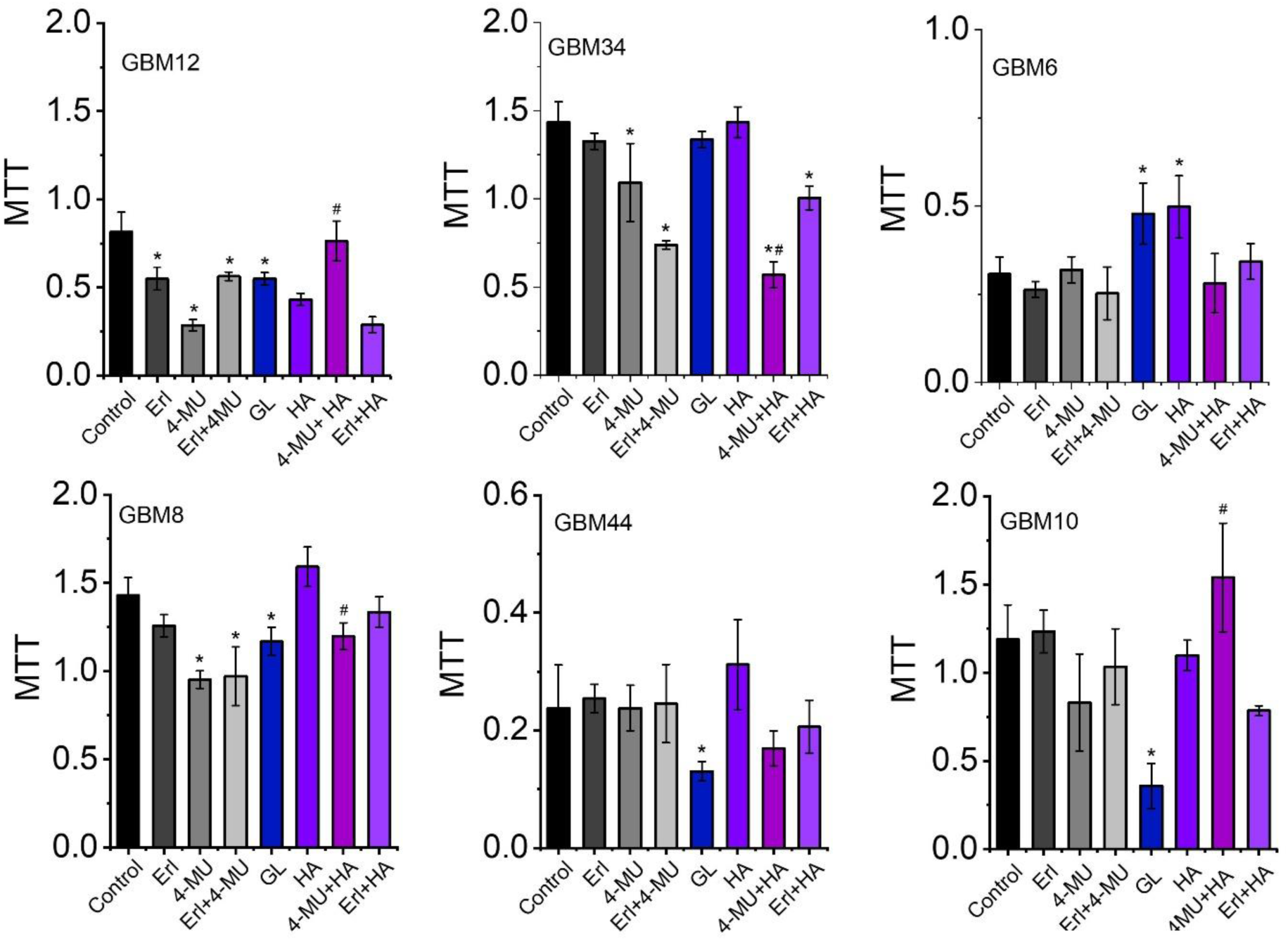
Low molecular weight hyaluronan selectively rescues metabolic activity of glioblastoma cells after treatment with inhibitors that target HA production. MTT cell viability assay (CyQUANT, Thermo Fisher Sci.) of GBM tumor cells exposed to different treatments. Inhibition of HAS preferentially affects GBM12 and GBM8, both show MGMT methylation status. Erlotinib and 4-MU show synergistic effect in GBM34. Metabolic activity is rescued by exogeneous LMW HA in GBM12, GBM8 and GBM10. HYAL inhibitor GL decreases proliferation of GBM12, GBM8, GBM44 and GBM10, contrary to GBM6. LMW HA ∼5 kDa 2.5mg/ml; *p < 0.01 (control), #p < 0.01 (4-MU).

Subsequently, GBM 3D cultures were incubated with LMW HA (∼5 kDa). Upon this treatment we observed a significant rescue of metabolic activity in GBM12, GBM10 and GBM8. Both primary tumors (GBM8 and GBM12) show O6-Methylguanine-DNA Methyltransferase (MGMT) promoter methylation. Soluble LMW HA increases cell metabolic activity in GBM6 and GBM8. These GBM tumors belong to the classical molecular subtype, characterized by EGFR overexpression and absence of TP53 deletion. Despite the low number of tumors analyzed here, these studies suggest that the rescue of cell activity is dependent on the tumor subtype, and in this case relates with their respective methylation profiles. Previous studies have demonstrated that the oncoprotein EGFRvIII sensitizes some subtypes of GBM tumors to current standard of care treatment by upregulating mismatch repair (MMR) proteins in MGMT methylated tumors (32). Hence, we are interested in investigating whether HA saccharide sequences may have a role in the sensitization or resistance mechanisms of GBM.

In healthy tissues most hyaluronan is of high molecular weight, however, tumors and tumor interstitial fluid, show elevated levels of low molecular weight (< 10kDa) HA (33,34). Hyaluronidases 1 and 2 (HYAL1 and HYAL2) have been identified as the main isoforms that degrade HA into small fragments. GL has been reported as an inhibitor of hyaluronidases (35). Our results show that GL reduces metabolic activity in all tumors, more significantly in GBM10 (**Figure 2**).

Hyaluronan oligosaccharides (< 5 kDa), contrary to larger chains, have been shown to selectively promote apoptosis in cancer cells as opposed to healthy cells (36). Low molecular weight HA has also been shown to increase sensitivity to chemotherapeutic drugs (37). In breast tumors, Hippo signaling pathway has been revealed as a potential antitumor treatment; high molecular weight hyaluronan demonstrated the maintenance of the Hippo signaling activation through CD44 interactions, upon degradation into smaller fragments, the cancer becomes more aggressive (38). Hence, we sought here to interpret the differential function of cell-secreted HA in GBM tumor progression and therapeutic resistance.

### Hyaluronan-regulated changes in GBM mitochondrial respiration

Due to the ubiquity of HA in the brain tissue and tumor microenvironment, several research groups have reported in recent years the role of hyaluronan as a nutrient in tumor progression (7,9,39). Moreover, research has revealed that mitochondrial DNA content is related to survival and that tumor cells activate compensatory pathways to maintain the mitochondrial metabolism (40,41), uncovering the possibility of promising combinatorial treatments.

We assessed mitochondrial respiration via a Seahorse Cell Mito Stress Test (Agilent) using Agilent Seahorse XFe96 extracellular flux analyzer and XFe96 spheroid microplates to evaluate the oxygen consumption of GBM cells within 3D hydrogel tumor tissues (GBM10, GBM12 and GBM34) (**Figure 3**). Our results show that GBM12 and GBM10 cells do not suffer a significant metabolic shift in the presence of HA. GBM10 shows a large maximal respiration capacity, while GBM12 cells appear more quiescent. GBM10 cells have a higher basal respiration rate than GBM12, also revealing a significant increase in the reserve capacity, non-mitochondrial respiration, and ATP-linked respiration (**Figure 3A**). Interestingly, the maximal respiration capacity in GBM34 cells increases with the presence of hyaluronan, further incentivized with MMW HA, suggesting that HA increases the capacity of these glioma cells to cope with the microenvironmental constraints (**Figure 3B-C**).

**Figure 3.**
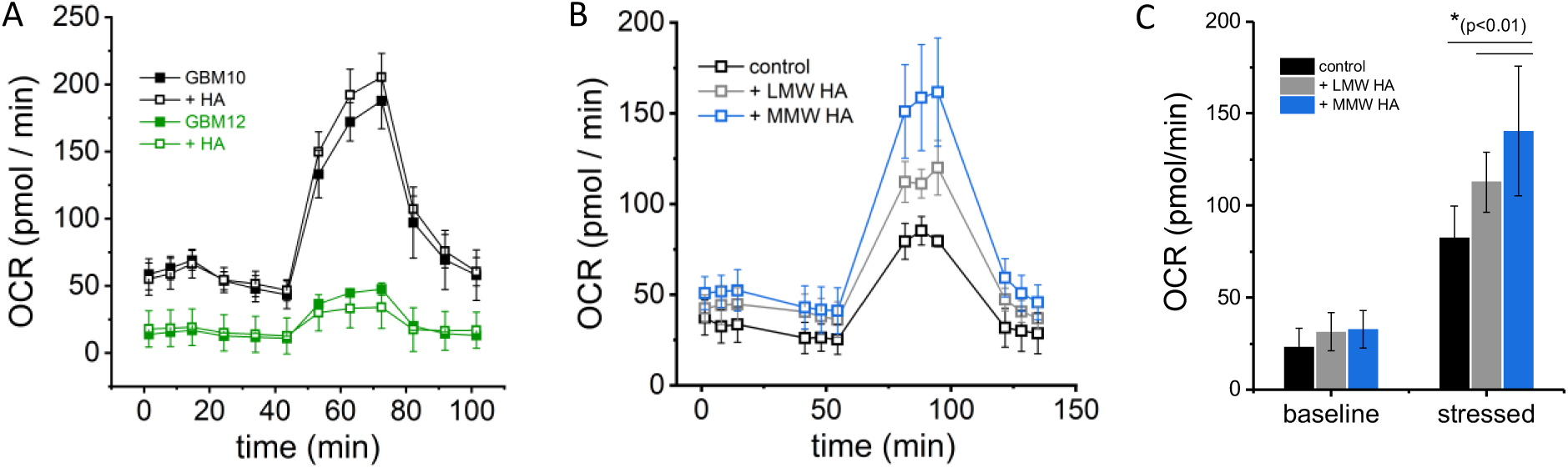
Mitochondrial activity dependence on exogeneous hyaluronan fragments. We used Seahorse XF technology (Agilent) to measure the oxygen consumption rate (OCR) over time (A-B), and the maximum OCR (B), in the medium immediately surrounding the GBM cells within 3D hydrogel platform, in a spheroid microplate. OCR is proportional to mitochondrial respiration. No significant difference was found in GBM10 and GBM12 with HA, were GBM12 shows a more quiescent phenotype than GBM10 (A). GBM34 shows increased mitochondrial respiration and metabolic potential in HA, more important in the presence of MMW HA (B-C). Solutions of HA 2.5mg/ml, density of 100k cells / gel/ well, 6 technical repetitions. *p<0.01

Metabolic reprogramming allows cancer cells to resist therapeutic interventions and become more invasive, showing a unique adaptability to changes in the microenvironment that include decrease oxygen and nutrient availability. Cancer cells are characterized by increased glycolysis, even in the presence of oxygen, in addition to the utilization of lipids, carbohydrates and amino acids as energy sources; however, studies have revealed subpopulations of cells with tumorigenic potential that rely on mitochondrial respiration versus glycolysis (41). A better understanding of the dynamic metabolic changes in GBM tumors can provide more efficient targeted interventions. In this work we exploit the distinct characteristics of 3D tumor models to analyze the role of hyaluronan in mitochondrial respiration. Notably, GBM12 cells displayed decreased proliferation capacity via exposure to LMW HA, while exhibiting more quiescence, contrary to GBM34 and GBM10 that show a higher metabolic activity and unresponsiveness to HA (**Figure 2**). While hyaluronan-regulated, more proliferative phenotypes, are associated to increased mitochondrial respiration, the HA-associated decrease in more quiescent cells do not correlate with alterations in oxygen consumption (**Figure 3**). These results illustrate the dual role of HA fragments in cell metabolism that can be used to increase mitochondrial function in GBM34 or to protect against apoptosis or increase proliferation by initiating alternative signaling pathways (GBM10). We show that these models can provide relevant information about GBM tumors metabolism. These findings can advance the understanding of the contribution of extracellular matrix components in tumor cell phenotypes to the regulation of mitochondrial function, leading to alternative therapeutic interventions.

### Hyaluronan inhibitors induce differential low molecular weight HA secretion in GBM cells

We isolated media samples from different GBM tumor samples after treatment with HAS, HYAL and EGFR inhibitors. After several purification steps we collected the hyaluronan and analyzed the molecular weight distribution with size exclusion chromatography. Results are shown in **Figure 4**. We observe that the differential expression across all samples lies in the region of 3-10 kDa, considered LMW HA. Notable differences are not seen after treatments in the high molecular weight region, while an increase of 35 kDa HA is observed after combinatorial treatment of erlotinib and 4MU in GBM8 and GBM10, and with GL in GBM44. Overall, we distinguish two groups of GBM types, the ones that do not express significant concentrations of LMW HA, i.e., GBM8 and GBM12, and the rest of GBM tumors that show a peak in the 3-10 kDa region. Among the latter, only GBM6, GBM34 and GBM44 show an enhanced secretion of LMW HA upon treatment with HYAL inhibitor GL. Both GBM8 and GBM12 are the only tumor samples in this study that show high MGMT promoter methylation status. Multiple studies have found that MGMT promoter methylation is associated with longer survival in diagnosed GBM patients that have been treated with temozolomide (TMZ). However, this is not a predictor factor for some patient cohorts (42,43).

**Figure 4.**
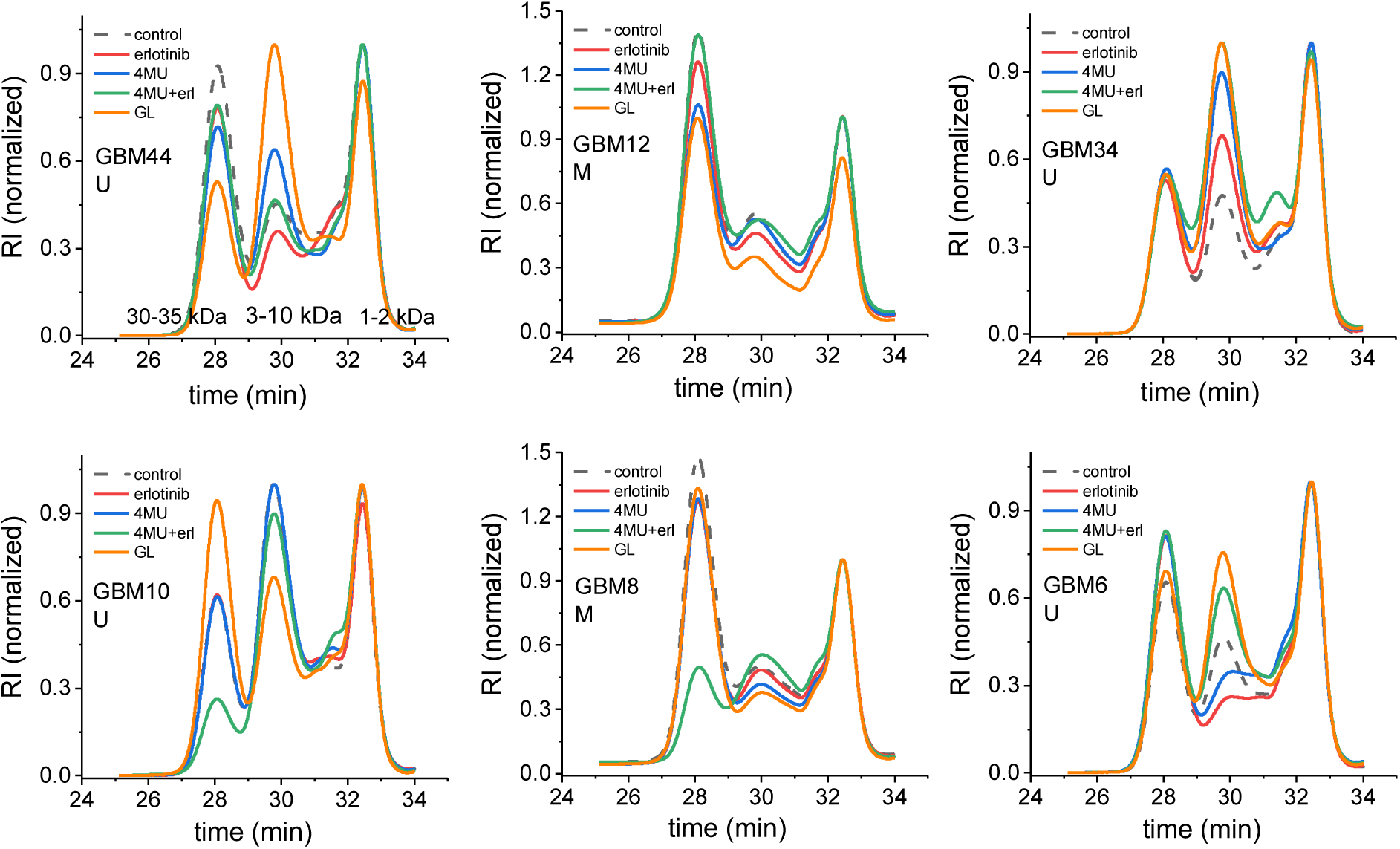
Hyaluronan inhibitors induce differential low MW HA secretion in GBM cells. Cell culture media was isolated from 3D cell cultures under the different treatments. Then, size exclusion chromatography (SEC) was used to separate and analyze the relative concentration of the molecular weight distribution of cell secreted hyaluronan fragments. MGMT status: U (Unmethylated), M (Methylated).

We investigated the effects of hyaluronan turnover in tumor progression and therapeutic response as a means to potentially identify alternative prediction factors. Both GBM6 and GBM8 exhibited increasing metabolic activity with supplementation of LMW HA (**Figure 2**), but they show different molecular weight distributions of secreted HA (**Figure 4**). Exogeneous HA can recuperate the metabolic activity of GBM10, GBM8 and GBM12, suggesting 4-MU efficacy may be dependent of HA availability in some GBM tumor subtypes. Those tumor specimens (GBM8 and 12) also display a lower production of LMW HA (**Figure 4**), suggesting the use of this ECM component to regulate their survival. HMW-HA is degraded by elevated hyaluronidases, and the interaction of LMW-HA with CD44 leads to pro-oncogenic cellular actions (44), however, the specific molecular mechanisms in these highly heterogeneous tumors remain unknown. Our results suggest that HA secretion profiling could be a useful tool to stratify tumors that will be susceptible to combinatorial treatments with HAS and HYAL inhibitors.

### Classical subtypes show an increase in HAS2 and HAS3 upon hyaluronidase inhibition

We subsequently investigated HAS and HYAL expression as a result of different treatments (4-MU and GL). HA is produced by the HA synthases, which can be inhibited in vitro by 4-MU (45). The silencing of HAS3 and blocking of CD44 have shown effective inhibition of glioma proliferation in vitro (46). We report here that HAS2 and HAS3 are selectively overexpressed with GL in GBM6, GBM8 and GBM12 while HAS1 is inhibited. The differential expression of the two classical subtypes (GBM6 and GBM8) is the most significant (**Figure 5 A-C**). The overexpression of epidermal growth factor receptor (EGFR) in these glioma tumors can indirectly decrease the expression of HAS, when the HA-mediated CD44-EGFR signaling pathway is inhibited. Moreover, HA fragments interact with the HA receptors differently depending on their MW (38). Several studies have shown that the addition of small HA fragments can disrupt the HA-CD44 interactions that results in the increased cancer aggressiveness (33).

**Figure 5.**
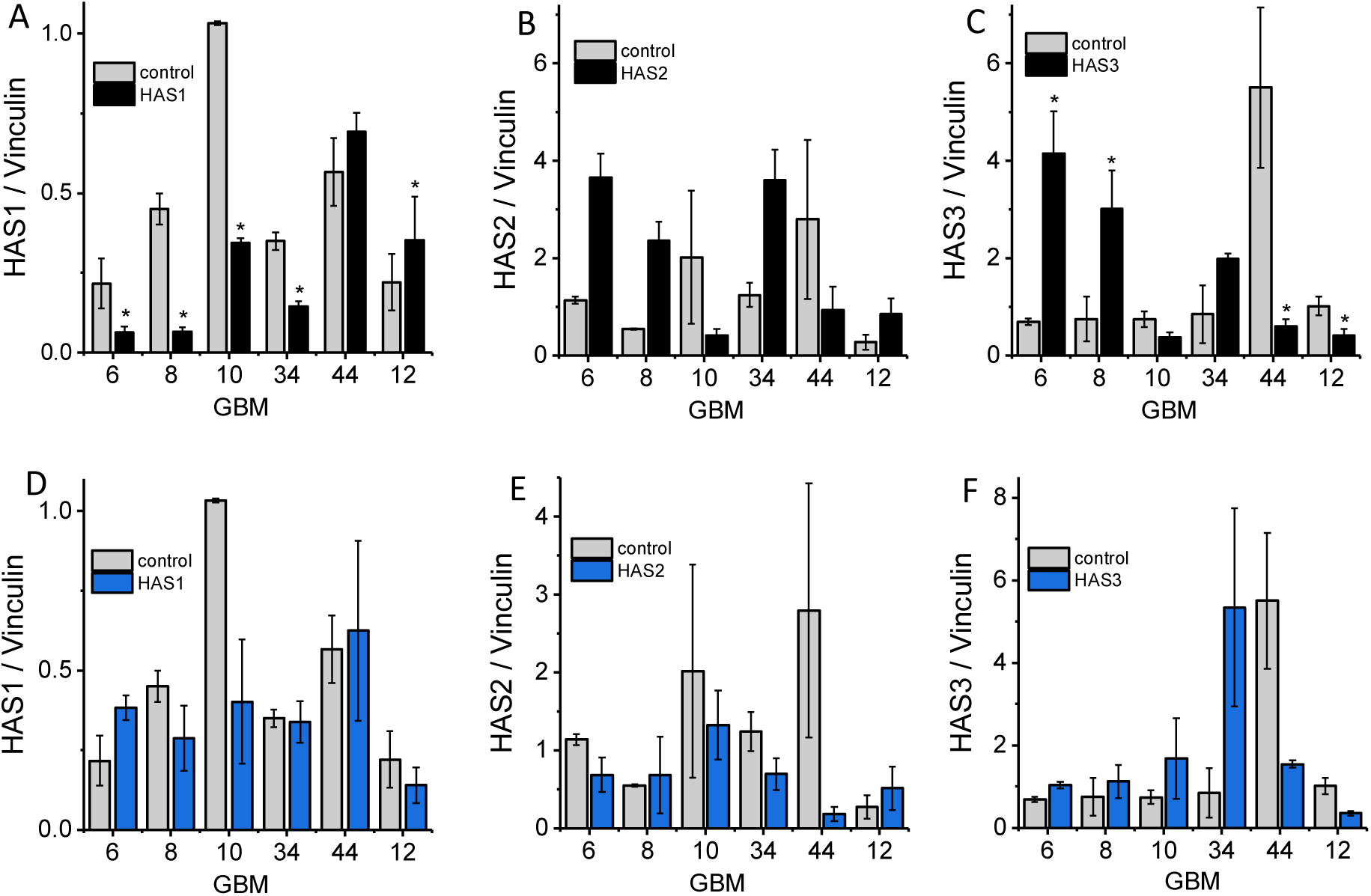
Classical subtypes increase HAS2 and HAS3 expression upon HA synthesis inhibition. Western Blot analysis of GBM platforms show differential expression of HAS1, HAS2 and HAS3 after treatment with (A-C) HYAL inhibitor (GL) and (D-F) HAS inhibitors (4-MU). Vinculin is used as a loading control. *p<0.05.

We analyzed HA-related synthase expression to understand the most relevant in response to constraints in the tumor microenvironment (**Figure 5, Figure S2**). In our group of tumor samples, we observe 4-MU selectively inhibits HAS depending on the tumor type. Upon treatment with 4-MU, HAS3 and HAS1 are inhibited in the mesenchymal subtypes while HAS2 is inhibited in the classical and proneural. Inhibition of hyaluronidase also triggers the upregulation of HAS in different manners depending on the tumor type. Mesenchymal subtypes (GBM10, GBM12, GBM44) tend to overexpress HAS1 whereas the remainder (classical and proneural) upregulate HAS2 and HAS3. HAS3 and receptor CD44 have been shown to be expressed at high levels in human glioma tissues and negatively correlated with the prognosis of patients with glioblastoma (46).

4-MU operates as a competitive substrate for UDP-glucuronyltransferase, an enzyme involved in HA synthesis. As a result, 4-MU treatment has been shown to lower HAS expression, thus reducing the production of HA (47). Glycyrrhizin from liquorice root acts as a potent inhibitor of HYALs in vitro (35), and the inhibition of HYAL has shown some therapeutic potential in certain cancers. The combination of standard of care GBM therapeutics and HAS and HYAL inhibitors may be a potent alternative to current therapeutic approaches.

### Glioma cells produce different cellular configurations of HA in response to inhibitors

The elevated production of HA without fragmentation has been associated to cancer progression (10,48). Increased deposition of hyaluronic acid is also a known characteristic of glioblastoma and other solid tumors (46,49). We observed a clear difference between the exogeneous deposition of HA in GBM6 and the cell-localized deposition of HA in most of the subtypes (endogenous matrix) (**Figure 6, Figure S3**). We observed cable-like structures of HA in GBM6 upon the treatment with erlotinib and GL. These structures originate from the cell surface and bind several cells together in a tubular-shaped network. These structures have been observed in other inflammatory conditions, where these HA cable-shaped structures stimulated the adhesion of leukocytes, without altering HA secretion (50). Brain tumors can establish communicating networks, long functional protrusions that increase tumor invasion and therapeutic resistance (51). The idea that an ECM component can assist in the effective communication of tumor cells opens the door to alternative treatments.

**Figure 6.**
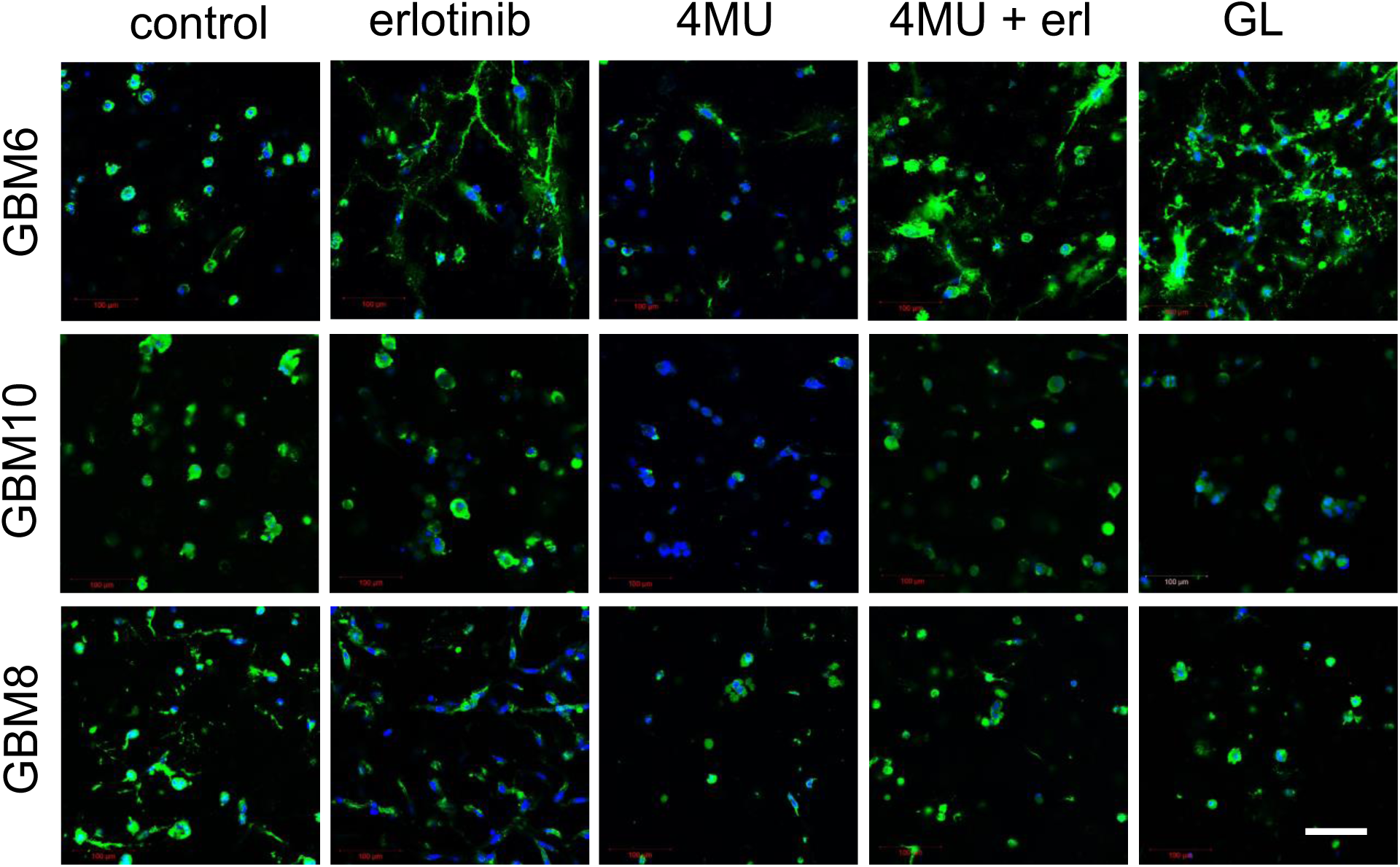
Different cellular configuration of HA in response to inhibitors. Hyaluronan in GBM tumor samples is identified with biotinylated HA binging protein (HBP) conjugated with Streptavidin-Alexa Fluor^®^ 488 (green) and DAPI for cell nuclei (blue). Scale bar 100 μm. HA affects metabolism

We propose here a model that considers the influence of hyaluronan in both PI3K signaling dependent and independent pathways, depending on the tumor molecular signatures. We have previously used these 3D models to prove that hyaluronan-dependent EGFR–CD44 interactions regulate phosphatidylinositol 3-kinase (PI3K) activity, that activate cell-survival and proliferation signaling, that also directly relates to HA secretion (14). PI3K signaling pathways are inhibited by 4-MU, erlotinib and GL in those tumors with depleted LMW HA production (GBM8 and 12) (**Figure 7A**), decreasing metabolic activity. Therefore, the addition of hyaluronan oligomers (**Figure 7B**) may rescue cell activity. However, in a subgroup of these cell tumors (i.e., GBM6) with constitutively active mutant of EGFR (EGFRvIII), inhibitors are not efficiently decreasing cell proliferation and HA accumulation can lead to increased glycolysis and changes in cell metabolism that support cell progression (**Figure 7C**).

**Figure 7.**
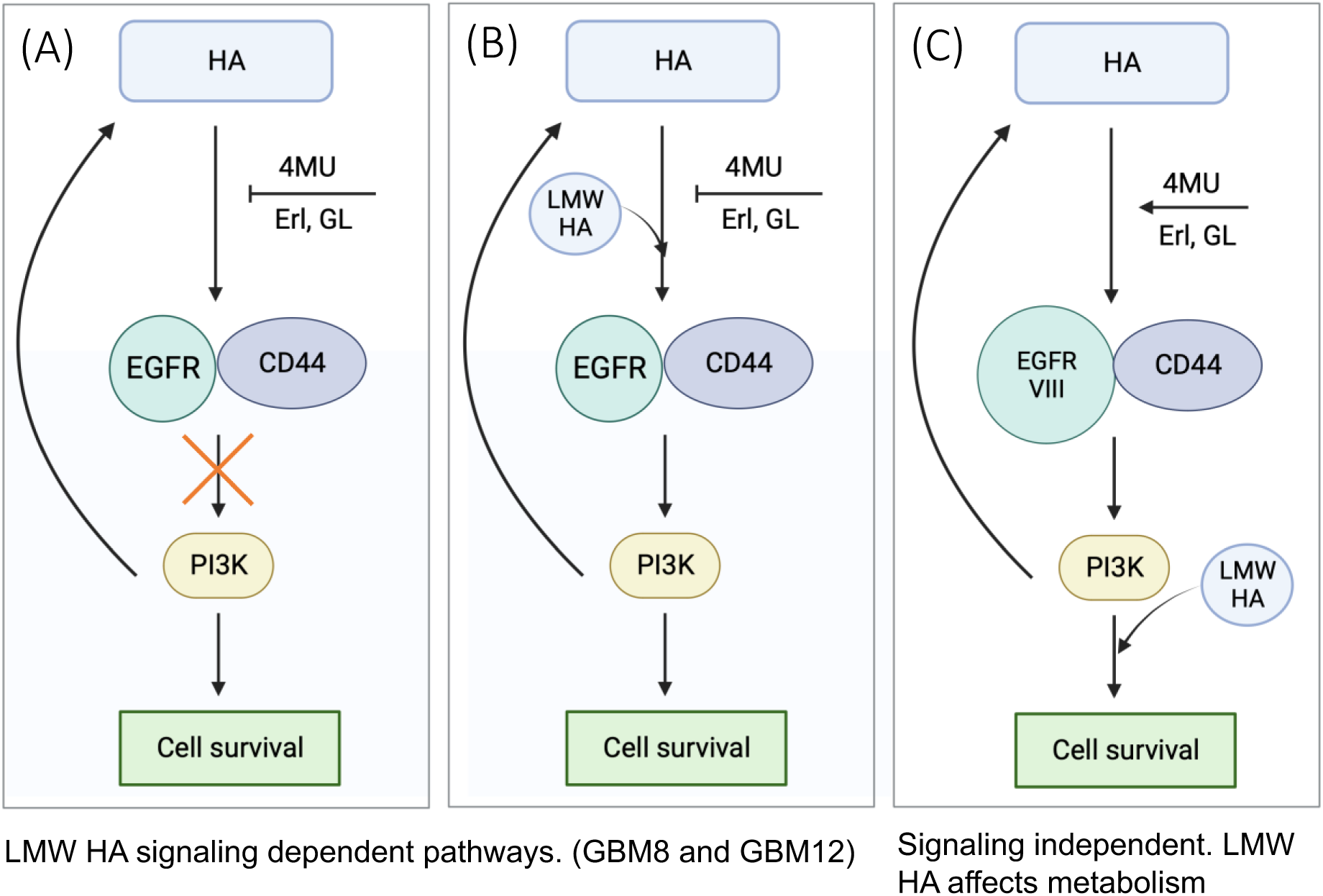
Hyaluronan EGFR-CD44-PI3K signaling dependent and independent GBM cell survival. Hyaluronan-dependent EGFR–CD44 interactions regulate phosphatidylinositol 3-kinase (PI3K) activity to initiate cell-survival and proliferation signaling, that also directly relates to HA secretion. These pathways are inhibited by 4-MU, erlotinib and GL in those tumors with low LMW HA production (GBM8 and 12) (A). Therefore, the addition of hyaluronan oligomers rescues cell activity (B). However, in tumors with EGFRvIII mutations (GBM6), with PI3K pathways constantly active, inhibitors are not efficiently decreasing cell proliferation, in addition, HA accumulation leads to increase glycolysis and changes in cell metabolism (C).

## Conclusions

Several in vitro and in vivo studies have described the capacity of brain tumors to express a spectrum of HA saccharides; they also express hyaluronidases that can degrade extracellular hyaluronan into low molecular weight HA, producing a tumor microenvironment governed by HA related interactions. GBM tumor cells produce HA of different molecular weights and are also influenced by exogenous HA produced by the stromal cells. The understanding of the subsequent activation of paracrine interactions towards biomass production and signaling pathways via ligand-receptor interactions (CD44-EGFR) are important in cancer growth and the capacity to discover new therapeutic targets in glioblastoma care. This work reveals that low molecular weight HA confers some glioblastoma tumors with the capacity to recuperate from the effects of targeted inhibitors, and that HYAL inhibitors can sensitize tumor cells by selectively inducing the accumulation of hyaluronan. This preclinical tool can provide data to predict the outcome of more precise combinatorial treatments in glioblastoma.

## Acknowledgements

We would like to acknowledge the following institutes for access to their facilities and services: the Roy J. Carver Biotechnology Center at UIUC, the School of Chemical Sciences Microanalysis Laboratory, the Tumor Engineering and Phenotyping Core at the Cancer Center at Illinois, the Carle R Woese Institute for Genomic Biology and the Materials Research Laboratory. Research reported in this publication was supported by the National Cancer Institute of the National Institutes of Health under Award Number R01 CA256481 (BACH) as well as the Bahl Family Research Fund from the Cancer Center at Illinois (BACH). The interpretations and conclusions presented are those of the authors and are not necessarily endorsed by the National Institutes of Health. We thank the Cancer Center at Illinois at the University of Illinois Urbana-Champaign for supporting this work. This project is supported by funding made available through the Cancer Center at Illinois Pilot Grant Program.

## Supplementary Material

**Figure S1.**
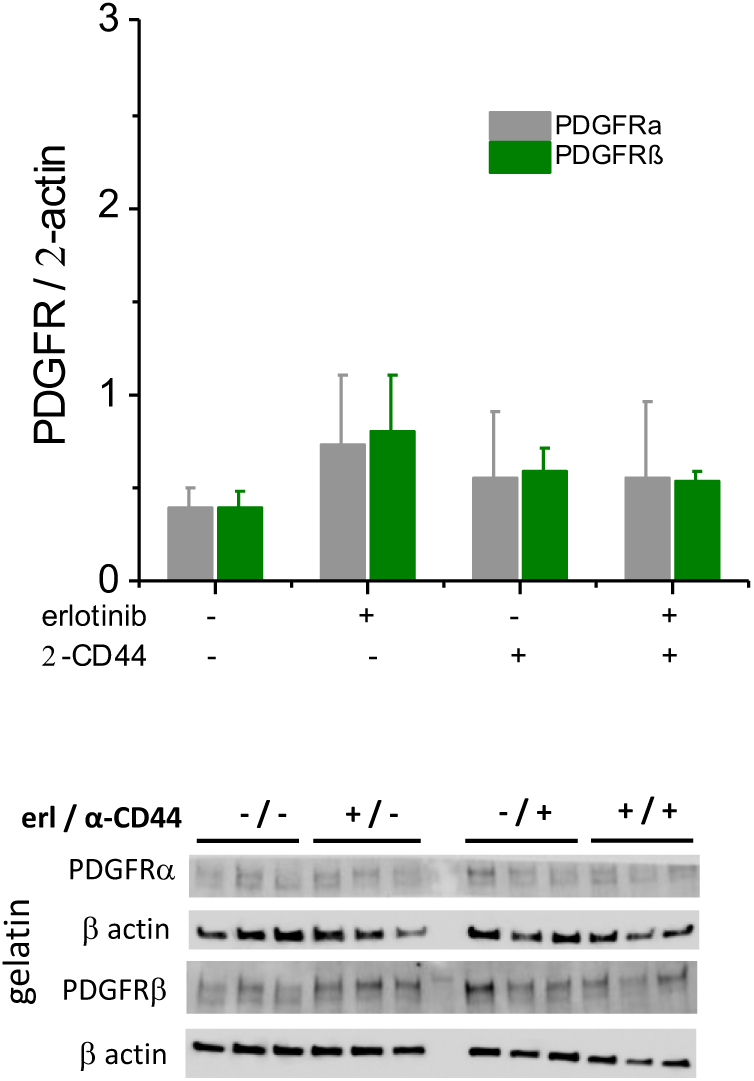
Western Blot analyses of PDGFR expression of GBM10 cells in GelMA-only hydrogels.

**Figure S2A.**
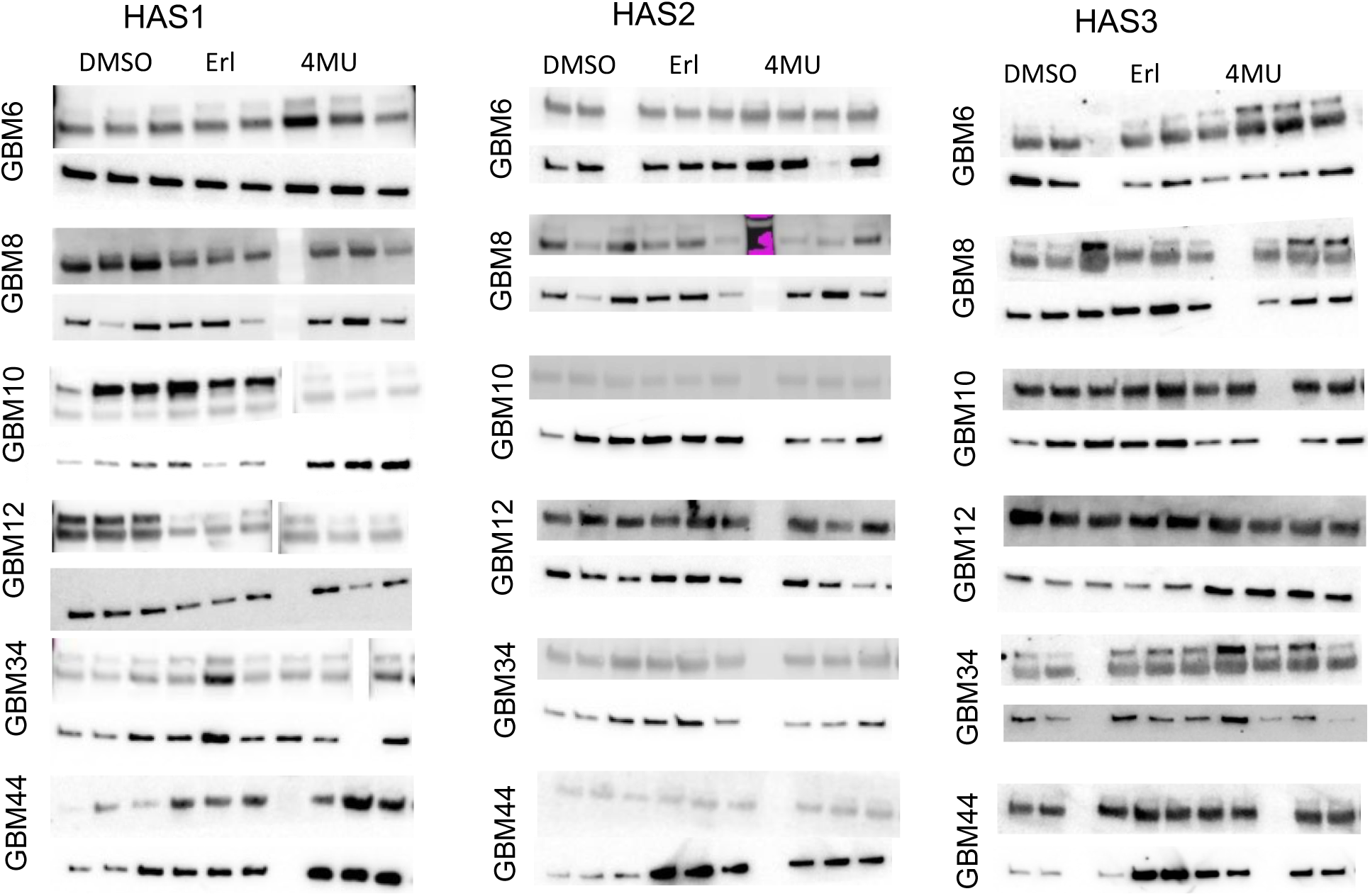
Western Blot analysis of GBM platforms show differential expression of HAS1, HAS2 and HAS3 as exposed to different inhibitors (erlotinib and 4-MU). Vinculin is used as the loading control.

**Figure S2B.**
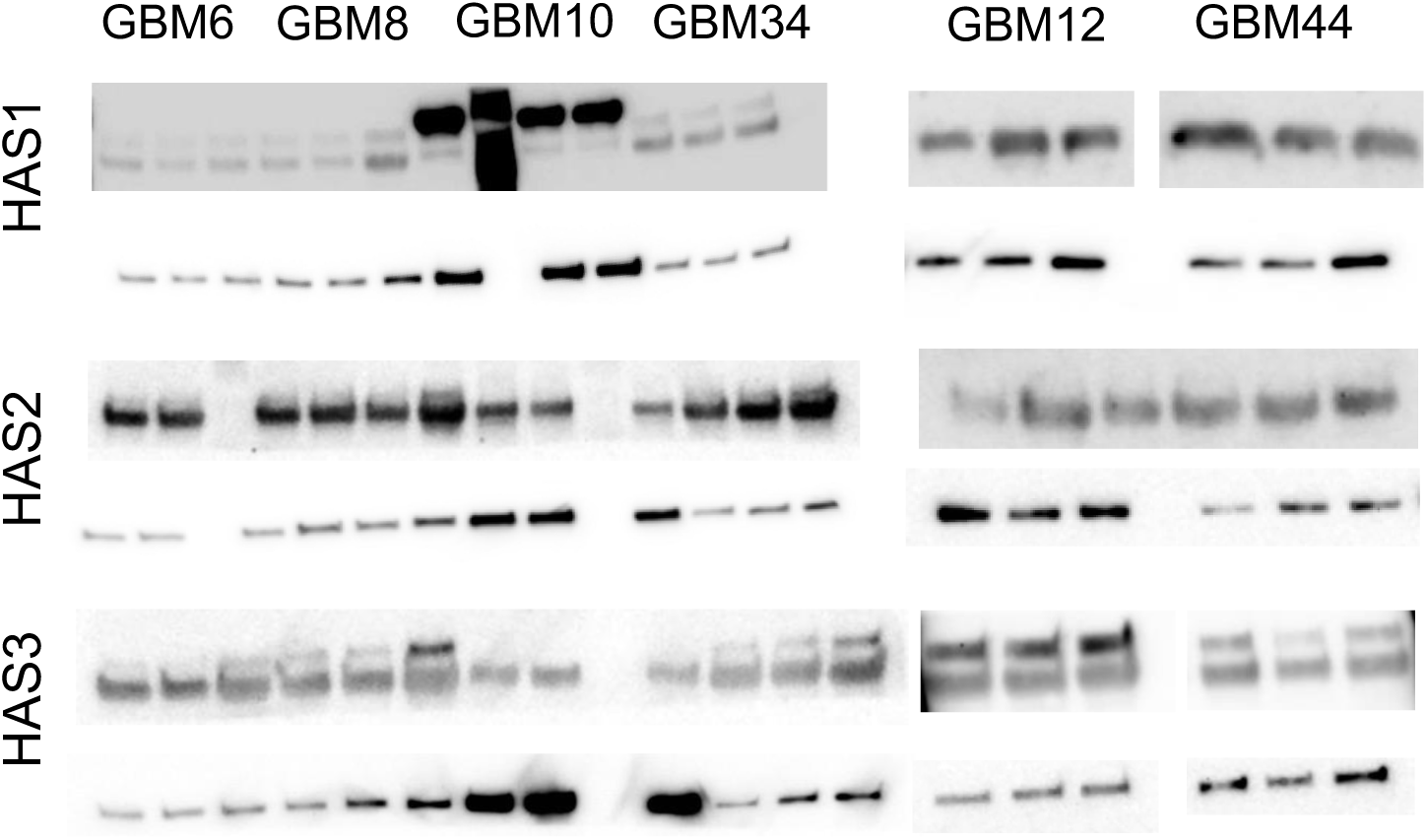
Western Blot analysis of GBM platforms show differential expression of HAS1, HAS2 and HAS3 as exposed to different inhibitors (GL). Vinculin is used as the loading control.

**Figure S3.**
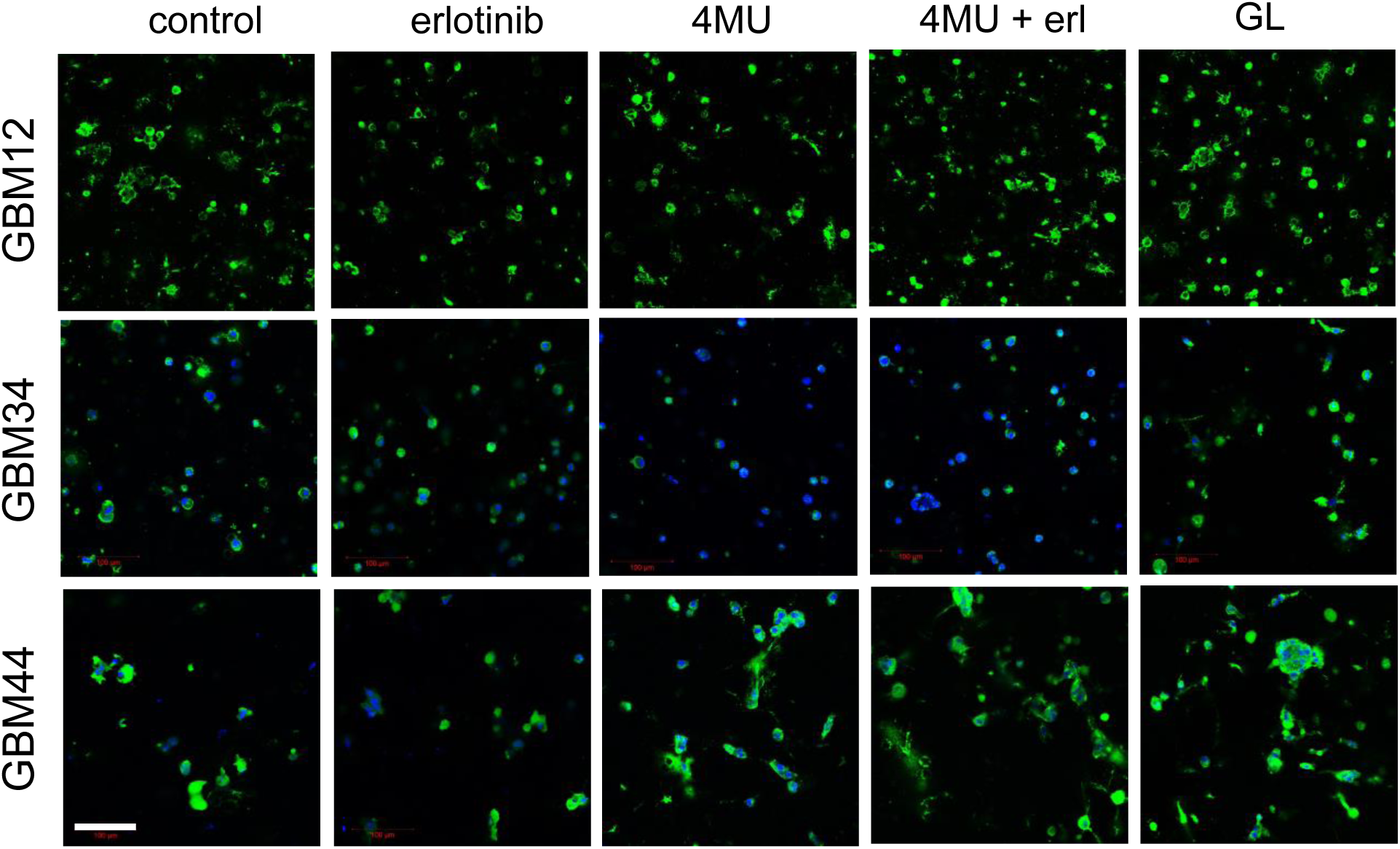
Different cellular configuration of HA in response to inhibitors. Hyaluronan in GBM tumor samples is identified with biotinylated HA binging protein (HBP) conjugated with Streptavidin-Alexa Fluor^®^ 488 (green) and DAPI for cell nuclei (blue). Scale bar 100 μm.

